# Longitudinal treatment selection under irreversible discontinuation: History-aware prediction and trajectory-aware learning in peanut-allergy prevention

**DOI:** 10.64898/2026.10.09.757908

**Authors:** Ronen Taub, Tanya Wasserman, Yonatan Savir

## Abstract

Clinical decision-support systems that guide longitudinal treatment must determine not only which treatment is preferred, but also when the available evidence is sufficient to change course. This challenge is especially important when discontinuation is irreversible, because stopping too early prevents later information from informing care. Food allergy is a growing public-health concern that can cause severe reactions and imposes substantial medical, dietary, and psychosocial burdens. Prevention is therefore a major clinical priority. In peanut-allergy prevention, this longitudinal decision is particularly consequential because resuming peanut consumption after discontinuation may be unsafe without renewed clinical assessment. In this work, we introduce a framework for preferred-treatment prediction and evaluate it using data from 628 children assigned to early peanut consumption or avoidance in the Learning Early About Peanut Allergy trial. We first introduced visit-wise and history-aware models that update recommendations as clinical and immunological measurements accumulate. Preferred treatment was poorly discriminated at baseline but became substantially more predictable during follow-up, with history-aware models achieving area-under-the-curve values of about 0.9 at later visits. However, imposing irreversible discontinuation after training significantly reduce the performance of these models. We therefore developed TreatNet, which jointly optimizes visit-specific predictors through a trajectory-level objective representing the downstream consequences of early discontinuation. Under the same irreversible constraint, TreatNet achieved accuracies of 0.805–0.839 and area-under-the-curve values of 0.819–0.869 across follow-up visits and two alternative preferred-treatment definitions. TreatNet generally preserved consumption while baseline evidence was limited and selectively recommended discontinuation after more informative measurements became available. These findings show that longitudinal information is most useful when model training reflects how current actions restrict future options. Although evaluated in peanut-allergy prevention, TreatNet provides a general framework for decision-support problems involving irreversible or difficult-to-reverse treatment changes.

**Author summary:** Food allergy can cause severe reactions and places substantial medical, dietary, and emotional burdens on children and their families. Early peanut consumption can prevent peanut allergy, but whether an individual child should continue or stop consumption may become clearer only as new clinical information is collected. This creates an important challenge: stopping too early may be difficult to reverse and can prevent later information from guiding care. We used data from 628 children in a major peanut-allergy prevention trial to study how recommendations can be updated over time. We first developed models that reconsider the preferred treatment at each visit, including models that use the child’s complete measurement history. We found that the preferred treatment was difficult to identify at baseline but became substantially more predictable during follow-up. However, models trained to make each decision separately lost much of this benefit when stopping was treated as irreversible. We therefore developed TreatNet, which learns how an early decision affects the choices available later. TreatNet generally continued consumption while evidence was limited and recommended stopping selectively after more informative measurements became available. Although tested in peanut-allergy prevention, this approach could support many healthcare decisions in which stopping or changing treatment restricts future options.

## Introduction

Food allergy is a growing public health concern, affecting an estimated 6–8% of children in industrialized countries [1]. Peanut allergy is among its most persistent forms and is associated with a substantial risk of severe reactions and anaphylaxis [1,2]. Beyond the risk of acute events, food allergy imposes considerable psychosocial and economic burdens through dietary restrictions, continuous monitoring, emergency preparedness, and healthcare use [3, 4]. Preventing peanut allergy during infancy has therefore become an important clinical priority.

The Learning Early About Peanut Allergy (LEAP) trial established that sustained peanut consumption from infancy through five years of age substantially reduces the development of peanut allergy in high-risk children compared with avoidance [5]. Follow-up studies showed that this protective effect largely persisted after a subsequent period of avoidance and into adolescence [6,7]. These findings transformed clinical guidance by supporting early peanut introduction for infants at elevated risk [8–10].

Nevertheless, the optimal management strategy for an individual child may not be fully apparent at baseline. Clinical and immunological measurements, including peanut-specific immunoglobulins, component-resolved markers, and skin-prick-test responses, change as the immune system develops [11–19]. Their predictive relevance also changes over time. Studies using LEAP and other cohorts have shown that oral food challenge outcomes can be predicted with increasing accuracy from measurements obtained later in childhood [13, 14, 20–22]. Thus, an uncertain baseline assessment may become substantially more informative after additional clinical observations have accumulated.

Most existing approaches do not make use of this temporal structure when selecting treatment. Current guidelines primarily apply rule-based baseline risk stratification to determine whether and how peanut should be introduced [8, 9]. Subsequent studies have refined this initial decision by identifying characteristics associated with peanut tolerance, estimating the benefit of early introduction across clinical risk groups, and deriving individualized treatment rules [23–27]. Although some analyses have included measurements obtained after baseline, they have generally estimated risk, treatment benefit, or a treatment recommendation at a particular decision point. They have not treated the child’s evolving measurements as a sequence of interdependent decisions in which a recommendation can be reconsidered as additional evidence becomes available.

The first objective of this study was therefore to formulate peanut-allergy prevention as a longitudinal treatment-selection problem (Fig. 1). In a static strategy, the baseline recommendation is retained throughout follow-up, even though baseline information may be insufficient and more informative measurements subsequently become available. An adaptive strategy instead reassesses treatment at each visit using the information available at that clinical time point. This distinction separates the question of whether an outcome can be predicted from the question of when the available evidence becomes sufficiently informative to support an individualized treatment recommendation. Longitudinal reassessment may allow treatment to be preserved while uncertainty is high and discontinued when later measurements indicate that continued intervention is unlikely to be necessary.

**Fig 1.**
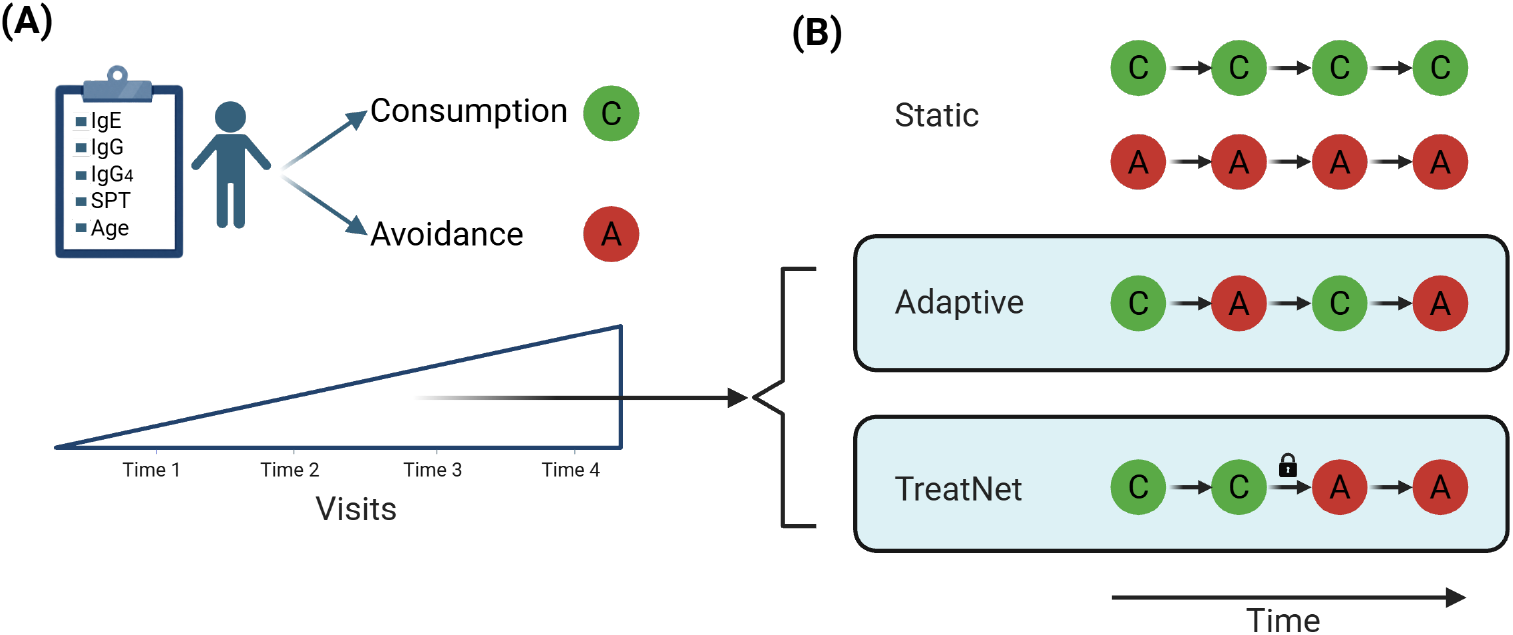
Longitudinal treatment strategy selection. (A) Peanut-allergy risk is only partially resolved at baseline, while clinical and immunologic measurements evolve across visits. These temporal dynamics motivate a dynamic treatment strategy in which recommendations are reassessed as the patient’s trajectory becomes clearer. (B) A static strategy retains the baseline recommendation and therefore cannot use information acquired during follow-up. We first introduce an adaptive strategy that updates the recommendation at each visit as the child’s clinical trajectory becomes clearer. However, because its visit-specific predictors are optimized independently, this strategy may produce clinically inadmissible stop–restart sequences. We then introduce TreatNet, which extends adaptive treatment selection to settings in which discontinuation is effectively irreversible by jointly optimizing visit-specific predictors over the complete decision pathway. Treatment may continue while evidence accumulates or be discontinued once, after which it cannot be resumed. At each visit, predictions use only the information available at that clinical time point.

Repeated reassessment alone, however, does not ensure a clinically coherent treatment pathway. Predictors optimized independently at successive visits may generate stop–restart sequences, even when treatment cannot realistically be resumed after discontinuation. This issue is especially important when an early decision changes which actions remain available later: an incorrect early discontinuation cannot be corrected by a more accurate recommendation at a subsequent visit. Imposing an irreversible-stopping rule only after training may therefore eliminate much of the apparent benefit of adaptive prediction.

Sequential decision-making has been studied more broadly through dynamic treatment regimes and reinforcement-learning methods [28–32]. These approaches typically estimate the consequences of alternative actions from longitudinal variation in treatments assigned at successive decision points. LEAP, however, randomized treatment only once, at baseline. Participants predominantly followed either sustained peanut consumption or sustained avoidance, providing the two extreme treatment trajectories but not the intermediate stop–continue sequences required to estimate the causal effects of discontinuation at different visits. Consequently, conventional dynamic-treatment-regime methods cannot be applied without extrapolating outcomes for treatment sequences that were not observed.

The second objective of this study was therefore to develop a trajectory-aware learning framework for settings in which treatment is repeatedly reconsidered but discontinuation is effectively irreversible (Fig. 1). We introduce TreatNet, which jointly optimizes visit-specific predictors over the complete decision pathway. Treatment may continue while evidence accumulates or be discontinued once, after which it cannot be resumed. By representing this dependency during training, TreatNet accounts for the downstream consequences of early decisions rather than treating each visit as an independent classification problem. At inference, each predictor uses only the measurements available at the corresponding clinical time point; trajectory awareness arises from the learning objective, not from access to future information.

Using longitudinal data from LEAP, we first examined how the information available for predicting the final oral food challenge outcome changes across visits. We then defined retrospective preferred-treatment labels from the randomized treatment assignment and final challenge outcome and evaluated two alternative definitions for the subgroup whose preferred treatment remains uncertain. These labels provide a benchmark for comparing longitudinal prediction strategies rather than an estimate of each child’s causal optimal treatment, because outcomes under the unassigned treatment are not observed. We compared static strategies, independently optimized adaptive models, and TreatNet under the same irreversible-discontinuation constraint. Finally, we analyzed individual prediction trajectories to determine how TreatNet balances uncertainty at baseline against the opportunity to discontinue treatment after more informative follow-up measurements become available.

## Materials and methods

### Study cohort

The Learning Early About Peanut Allergy (LEAP) clinical trial followed 628 infants randomly assigned to either peanut consumption or peanut avoidance. Participants were monitored across four clinical visits from infancy through early childhood. At each visit, a set of measurements was collected. For the present analysis, we use demographic attributes (age, ethnicity, sex), blood-based immunological markers (IgE, IgG, IgG_4_), and skin-prick-test wheal measurements. Together with the randomized treatment assignment and the oral food challenge (OFC) outcome, these repeated assessments provide a longitudinal view of patient trajectories under different treatment regimens.

The distribution of patient subgroups is shown in Table 1. A total of 54 patients failed the OFC while assigned to avoidance, leaving uncertainty as to whether early consumption might have altered their outcome. The next subgroup consisted of 260 patients (41.40%) who passed the OFC despite avoidance, suggesting that treatment may not have been necessary. Ten patients (1.59%) failed the OFC despite peanut consumption, indicating that treatment did not prevent allergy development and that exposure may have carried potential risk. Finally, 304 patients (48.41%) consumed peanuts and passed the OFC.

**Table 1.** Distribution of OFC outcomes across treatment groups.

| Treatment group | OFC outcome | Count | Percentage |
| --- | --- | --- | --- |
| Control | Failed OFC | 54 | 8.60% |
| Control | Passed OFC | 260 | 41.40% |
| Treatment | Failed OFC | 10 | 1.59% |
| Treatment | Passed OFC | 304 | 48.41% |

### Temporal evolution of predictive clinical information

We examined how the predictive signal evolves across visits. For this, we trained gradient-boosted decision tree model XGBoost [33] to predict the OFC outcome independently at each visit. XGBoost is well-suited for structured clinical data because it models nonlinear feature interactions and accommodates heterogeneous feature types.

After training, we interpret which clinical variables drive predictions at each time point using SHapley Additive exPlanations (SHAP) values [34]. SHAP provides a theoretically grounded feature attribution method based on cooperative game theory, quantifying each feature’s contribution to individual predictions. Aggregating SHAP values across patients yields visit-specific feature importance profiles that reveal how the relevance of clinical signals changes over time.

OFC prediction was evaluated using nested patient-level cross-validation. The outer cross-validation folds were used exclusively for held-out performance evaluation, whereas the inner cross-validation folds were used for XGBoost hyperparameter selection. Patient assignments were kept consistent across the four visits, such that the same outer-fold patients were evaluated at each time point. An independent XGBoost model was then trained for each visit using only the clinical information available at that visit. This design ensured that differences in predictive performance across visits reflected changes in the available clinical information rather than differences in the evaluated patient population.

SHAP values were computed on the held-out patients in each fold. Feature importance was quantified using the mean absolute SHAP value for each feature and subsequently aggregated across folds to characterize the importance and variability of clinical variables at each visit.

### Longitudinal treatment-selection framework

Having examined how the predictive information available for the final OFC outcome changes across visits, we next formulated the main treatment-decision problem considered in this study. This task is distinct from OFC prediction: rather than predicting whether a patient will ultimately pass or fail the OFC, the goal is to determine whether peanut consumption or avoidance is the preferred treatment at each visit as new clinical information becomes available. To define and evaluate this problem, we first establish retrospective preferred-treatment labels and compare static with longitudinally adaptive treatment strategies, thereby quantifying the value of reassessment as clinical information accumulates. We then evaluate whether this value is preserved under irreversible discontinuation by comparing independently optimized reference models with the trajectory-aware TreatNet framework.

### Preferred treatment definition

The LEAP dataset provides two signals that enable retrospective evaluation of treatment decisions: the final OFC outcome (pass or fail) and the original treatment assignment (consumption or avoidance). Using these signals, we define a notion of the *preferred treatment strategy in hindsight*. This definition does not attempt to reconstruct the true counterfactual outcome, which cannot be observed, but instead establishes a conservative benchmark for comparing treatment policies. In Figure 2, we describe the preferred treatment in hindsight for each subgroup and the logic behind the choice.

**Fig 2.**
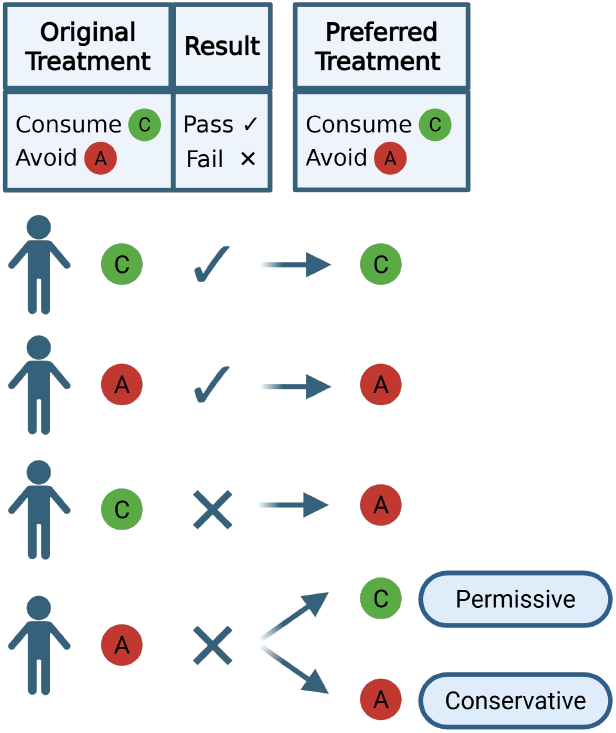
Definition of the preferred treatment in hindsight. The final OFC outcome and original LEAP treatment assignment were combined to define the preferred retrospective treatment label for each patient. For three outcome-treatment combinations, the preferred treatment can be concluded. For patients assigned to avoidance who subsequently failed the OFC, two alternative definitions were considered: a conservative profile, retaining avoidance as the preferred treatment, and a permissive profile, assigning consumption as the preferred treatment.

We distinguish between a conservative treatment approach and a permissive treatment approach. For three observed treatment–outcome combinations—avoidance/OFC-pass, consumption/OFC-pass, and consumption/OFC-fail, the retrospective decision rule assigns the same preferred, treatment label under both profiles. The remaining subgroup, comprising patients assigned to avoidance who subsequently failed the OFC, is inherently ambiguous because their outcome is undesired while the outcome under early peanut consumption was not observed. We therefore considered both possible labels for this subgroup. Under the conservative profile, avoidance remained the preferred label, whereas under the permissive profile, consumption was assigned as preferred. Each such profile creates a different set of labels, and as a consequence changes the label distribution greatly. Thus, we examined both to show the generality of our approach.

The resulting labels are used as a retrospective benchmark for learning when the observed patient trajectory supports continuation or discontinuation; they are not intended to estimate counterfactual biomarker trajectories following alternative earlier treatment decisions. Post-baseline predictions are based on the clinical trajectory actually observed under the participant’s assigned treatment. Accordingly, the analysis evaluates longitudinal treatment selection from observed information rather than the causal effect of hypothetical treatment switching at each visit.

#### Static, adaptive, and constrained adaptive policies

We formulated the clinical problem as a sequence of four treatment decisions corresponding to the LEAP study visits. We evaluated three treatment policies that differ in how recommendations are updated over time.

##### Static policy

A single treatment decision is made at baseline and retained throughout follow-up.

##### Fully adaptive policy

Treatment is reassessed independently at each visit using the information available at that time. This allows unrestricted transitions between consumption and avoidance across visits.

##### Constrained adaptive policy

Treatment may be continued or discontinued as new information becomes available, but once discontinuation is selected, treatment cannot subsequently be resumed. Thus, admissible trajectories contain at most one transition from consumption to avoidance.

#### Visit-wise adaptive treatment selection

We evaluated visit-wise adaptive treatment selection by comparing static strategies with models that update the preferred-treatment prediction at successive visits, Figure 3). Static strategies retain a single baseline recommendation, whereas visit-wise adaptive strategies use the clinical and immunological information available at each visit to revise that recommendation. We further distinguished between models that use only the current-visit measurements and models that use the complete measurement history available by that visit. These comparisons quantify the value of visit-wise adaptation as additional patient-specific information becomes available.

**Fig 3.**
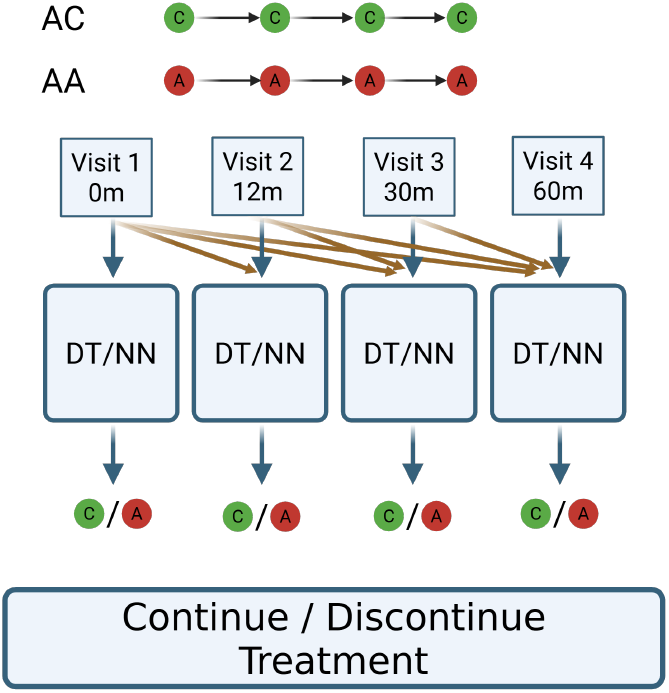
Visit-wise adaptive treatment selection. Static models make a single preferred-treatment prediction at baseline and retain it throughout follow-up, preventing them from using information acquired at later visits. Visit-wise adaptive models instead use an independent predictor at each clinical visit, allowing the recommendation to be updated as new clinical and immunological measurements become available. Standard visit-wise models use only the measurements collected at the current visit, whereas history-aware variants incorporate all measurements accumulated through that visit (red arrows).

##### Static models

We considered three time-invariant strategies that cannot use information acquired during follow-up. *Always Consume* assigns peanut consumption to all patients throughout follow-up, whereas *Always Avoid* assigns avoidance throughout follow-up. *Baseline MLP* uses a multilayer perceptron trained exclusively on baseline measurements to predict the preferred treatment, and its baseline prediction is retained across all subsequent visits.

##### Visit-wise adaptive models

We trained an independent predictor for each of the four visits, allowing the preferred-treatment prediction to be updated as new measurements became available. *Visit-wise DT* uses decision-tree-based models, whereas *Visit-wise MLP* uses multilayer perceptrons. Each visit-specific model receives only the measurements collected at the corresponding visit and is optimized independently. Differences in performance across visits therefore quantify how treatment selection benefits from the evolving clinical information available during follow-up.

##### History-aware adaptive models

We additionally evaluated *History-aware DT* and *History-aware MLP*. These models retain the visit-wise architecture but receive all measurements accumulated through the corresponding visit. Later predictors therefore incorporate progressively longer patient histories, allowing us to determine whether adaptive treatment selection benefits from the complete observed clinical trajectory beyond the information contained in the current assessment alone.

Together, these models establish whether visit-wise adaptive treatment selection improves performance relative to a fixed baseline strategy and whether accumulated clinical history provides additional benefit. However, each adaptive predictor is trained independently and optimizes performance only at its corresponding visit, without accounting for the consistency of the complete treatment trajectory. These models therefore also provide the principal reference setting for evaluating whether TreatNet can preserve the benefit of visit-wise adaptation when treatment discontinuation is irreversible.

Preferred-treatment models were evaluated using 10-fold patient-level cross-validation. Unlike the XGBoost analysis of OFC prediction, no hyperparameter search was performed for these models; the MLP architectures and decision-tree settings were fixed before evaluation. The same patient-level fold assignments were used across models and visits to enable direct comparison of treatment-selection strategies.

Treatment status was handled according to the information available in the intended clinical setting. At baseline, before a treatment decision has been made, randomized LEAP treatment assignment was excluded from the model inputs. At subsequent visits, current treatment status was included as a binary input together with the available clinical and immunological measurements, because the treatment currently being received is known when a continuation or discontinuation decision is made.

History-aware variants received the current-visit measurements together with all measurements observed at previous visits; at follow-up visits, current treatment status was included as an additional known contextual variable.

#### TreatNet: trajectory-aware optimization under irreversible discontinuation

TreatNet is designed to optimize treatment decisions over the complete longitudinal trajectory rather than at individual visits independently. Its architecture mirrors the visit-wise MLP reference model: one multilayer perceptron is assigned to each clinical visit, and each module predicts the preferred treatment using the information available at that time. A history-aware variant, *TreatNet (Hist*.), additionally provides each visit-specific module with all measurements accumulated up to that visit.

The distinction between TreatNet and the reference models lies in the training objective. In the visit-wise baselines, each predictor is optimized independently against the preferred-treatment label at its corresponding visit. Such training does not account for the downstream consequence of an early discontinuation decision. Under the constrained adaptive policy, however, once treatment is discontinued, subsequent treatment decisions are no longer available. An error at an early visit can therefore alter the entire remaining trajectory. TreatNet addresses this dependency by coupling the outputs of all visit-specific modules through a trajectory-level objective.

Let *p*_*t*_ ∈ [0, 1] denote the predicted probability of continuing treatment at visit *t*, for *t* = 1, …, *T*. Under an irreversible stopping policy, treatment can reach visit *t* only if continuation was selected at every preceding visit. We therefore define the probability of treatment remaining active through visit *t* as

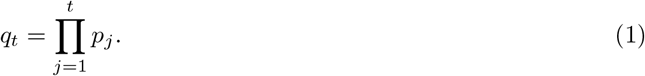

For four visits, the resulting trajectory terms are

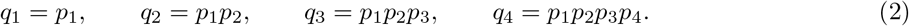

These cumulative products encode the path dependence introduced by irreversible discontinuation: a low continuation probability at an early visit suppresses the contribution of all subsequent visits, whereas later decisions remain relevant only when treatment has been maintained up to that point.

TreatNet combines these terms into a unified trajectory score,

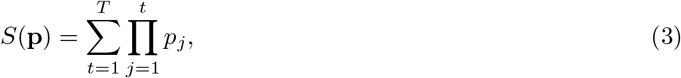

which for the four LEAP visits becomes

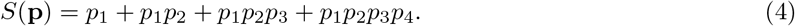

The optimization objective is then defined according to the retrospective preferred-treatment label,

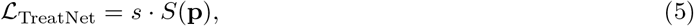

where *s* = +1 for patients whose preferred treatment is avoidance and *s* = *™*1 for patients whose preferred treatment is continued consumption. Minimizing this objective favors smaller cumulative continuation probabilities for patients assigned to avoidance and larger cumulative continuation probabilities for patients assigned to consumption. Because each term depends on all preceding visit outputs, gradients propagate across the trajectory rather than optimizing each visit in isolation.

This formulation directly reflects the clinical asymmetry of the decision problem. Continuing treatment preserves the option to reassess the patient at a later visit, whereas discontinuation removes that option. TreatNet therefore learns not only whether consumption or avoidance is preferred, but also how strongly an early decision should be weighted given its effect on all later decision opportunities (Figure 4).

**Fig 4.**
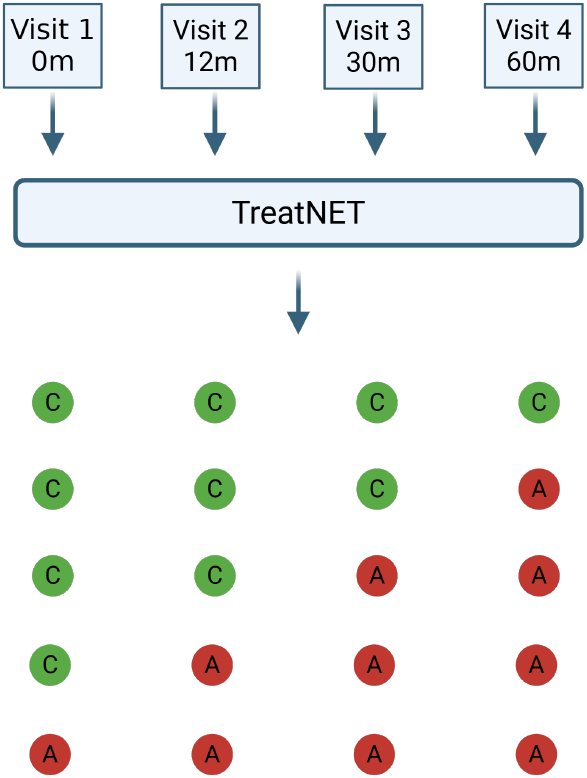
Trajectory-aware training of TreatNet. TreatNet contains one visit-specific predictor for each clinical time point, analogous to the visit-wise MLP reference model. Rather than optimizing the predictors independently, their outputs are coupled through a unified trajectory-level objective. The prediction at each visit contributes both directly and through cumulative products with the predictions of preceding visits, reflecting the irreversible stopping structure of the treatment policy. Consequently, early discontinuation decisions influence the contribution of all subsequent visits during training. At inference, TreatNet will favor patterns corresponding to the constrained adaptive policy. For the LEAP dataset, there are 5 such outputs (consumption advised at all visits, or discontinue at each of the visits).

The trajectory coupling is introduced during training only. At inference, each visit-specific module independently produces its treatment probability from the information available at that visit; no future measurements or future model outputs are used. Thus, TreatNet differs from the comparison models in how the predictors are jointly optimized, rather than through access to additional information or a different inference-time constraint.

### Evaluation under irreversible discontinuation

A single decision threshold was selected using the validation data and was shared across all visits. For unconstrained accuracy, visit-specific continuation scores were thresholded directly to obtain binary treatment recommendations. For constrained accuracy, the same threshold was first applied and the irreversible-discontinuation rule was then propagated through the trajectory: once avoidance was predicted, all subsequent visit predictions were set to avoidance. For unconstrained AUC, the original continuous continuation scores were used. For constrained AUC, the binary constrained trajectory was first obtained as described above and was then used to mask the original continuation scores. Thus, once the constrained trajectory entered the avoidance state, the corresponding continuation score and all subsequent continuation scores were set to zero. ROC AUC was calculated from these constraint-adjusted continuous scores.

### Model architectures

The visit-wise MLP consisted of four separate multilayer perceptrons, corresponding to the baseline, 12-month, 30-month, and 60-month assessments. Each network contained three fully connected hidden layers with 128, 64, and 32 units, respectively. Each hidden layer was followed by layer normalization, a Gaussian error linear unit (GELU) activation, and dropout with probability 0.2. A final fully connected layer produced a single logit, which was transformed by a sigmoid function to obtain a continuation score. The four networks had separate parameters and were trained independently to predict the preferred-treatment label at their corresponding visits. TreatNet used the same four-network MLP architecture. The distinction between TreatNet and the visit-wise MLP therefore arose from the optimization objective rather than model capacity: whereas the visit-wise MLP modules were optimized independently, TreatNet jointly optimized their outputs using the trajectory-aware objective described below. The visit-wise DT consisted of four independently trained decision-tree classifiers, one for each visit. Each tree used the Gini impurity criterion, a maximum depth of 3, a minimum of 40 samples required to split an internal node, and a minimum of 20 samples per leaf. Class weights were set using the balanced weighting scheme, and the random seed was fixed at 42. Each tree returned a class-1 score corresponding to treatment continuation. For history-aware variants, the model architecture and training procedure were unchanged. The input at each visit was augmented with all clinical and immunological measurements accumulated at preceding visits in addition to the measurements from the current visit.

## Results

### Temporal evolution of OFC predictive performance and clinical feature importance

We first examine how well the final oral food challenge (OFC) outcome could be predicted from the clinical information available at successive visits. For each visit, separate XGBoost models were trained using only measurements recorded at that visit. Hyperparameters were selected within the inner cross-validation folds, and performance was evaluated on patients held out in each of the 10 outer folds.

Discrimination improved progressively as longitudinal measurements became available (Fig. 5A). Mean ROC AUC increased from 0.614 at baseline to 0.691 at 12 months, 0.952 at 30 months, and 0.988 at 60 months. The increase in ROC AUC more clearly demonstrates the additional discriminative information provided by later measurements. As expected, performance was highest when using measurements obtained at the 60-month visit, contemporaneously with the final outcome assessment.

**Fig 5.**
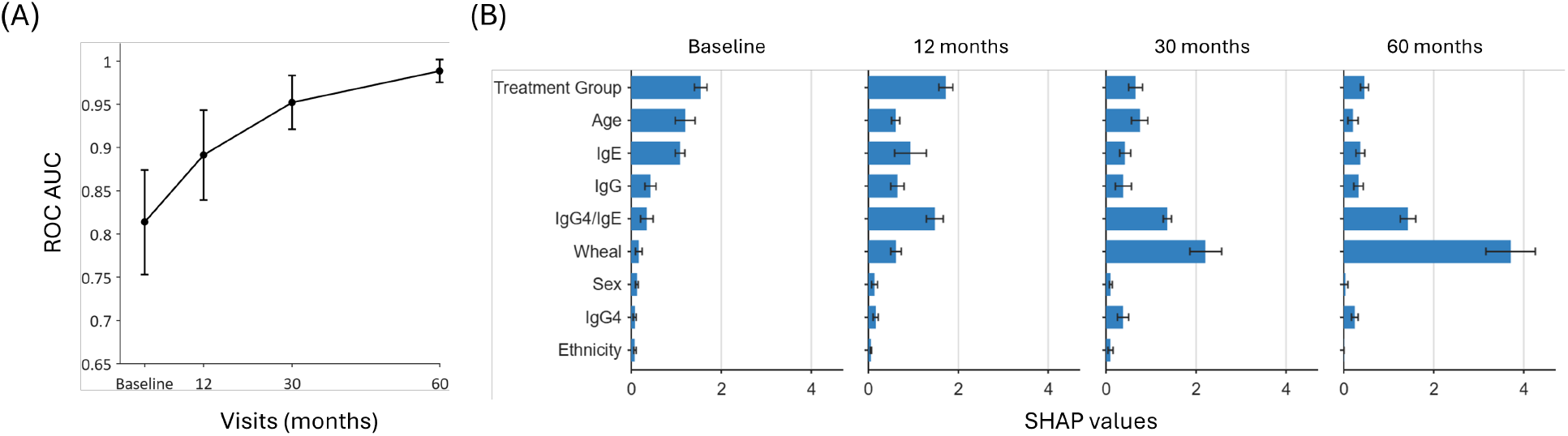
Temporal evolution of OFC predictive performance and clinical feature importance. (A) Separate XGBoost classifiers were trained at each clinical visit using only the measurements collected at that visit to predict the final OFC outcome. Points show the mean held-out ROC AUC across 10 outer patient-level cross-validation folds, and error bars denote 95% confidence intervals. Predictive discrimination increased progressively from baseline to the 60-month visit. (B) SHAP-based feature importance across visits. Bars show the mean importance rank across the outer folds, with lower ranks indicating greater importance; error bars denote standard deviations. Treatment assignment was the highest-ranked feature at baseline but declined in relative importance during follow-up, while the IgG4/IgE ratio and peanut skin-prick-test wheal size became the dominant predictors.

The SHAP analysis revealed that the variables contributing to prediction also changed over time (Fig. 5B). At baseline, treatment received was consistently ranked as the most influential feature, followed by the IgG4/IgE ratio and peanut-specific IgE. By 12 months, the IgG4/IgE ratio had become the highest-ranked predictor, followed by treatment received and peanut wheal size. At both 30 and 60 months, peanut wheal size was ranked first and the IgG4/IgE ratio second. Treatment received and peanut-specific IgE remained informative, but their relative contributions were lower and more variable across folds. In contrast, sex, ethnicity, and several individual immunoglobulin measurements generally occupied lower positions in the rankings.

These results demonstrate that the predictive representation of allergy outcome is temporally dependent. Baseline information provides moderate discrimination, but repeated clinical assessment progressively reveals signals that are more closely associated with the final OFC outcome. Moreover, the identity of the most informative measurements changes across visits, shifting from treatment received and baseline serological information toward the IgG4/IgE ratio and skin-prick-test response. Although SHAP importance reflects predictive association rather than causal effect, this temporal evolution supports the formulation of allergy management as an adaptive prediction problem in which treatment decisions can be reassessed as new patient-specific information becomes available.

### Visit-wise and history-aware adaptation improve preferred-treatment prediction

Having established that OFC predictive performance and clinical feature importance evolve across follow-up, we next examined whether the information acquired at successive visits improves prediction of the retrospectively preferred treatment. Static strategies retained a single baseline recommendation, whereas visit-wise adaptive models updated the recommendation independently at each visit. History-aware variants additionally incorporated all measurements accumulated through the corresponding visit. Performance was evaluated separately under the conservative and permissive preferred-treatment profiles (Table 2).

**Table 2.** Performance of static, visit-wise adaptive, and history-aware models for preferred-treatment prediction. Values show mean accuracy and AUC ± standard deviation across 10 outer patient-level cross-validation folds under the conservative and permissive preferred-treatment profiles. Static models make a single recommendation at baseline and retain it throughout follow-up; because their predictions do not change, they are reported only at baseline. Visit-wise adaptive models use measurements from the corresponding visit, whereas history-aware models use all measurements accumulated through that visit. Within each profile, the highest mean performance among the learned adaptive models at each follow-up visit is shown in bold.

| Visit | Metric | Always Avoid | Always Consume | Baseline MLP | Visit-wise DT | Visit-wise MLP | History-aware DT | History-aware MLP |
| --- | --- | --- | --- | --- | --- | --- | --- | --- |
| **Conservative profile** |  |  |  |  |  |  |  |  |
| Baseline | Accuracy | 0.522 ± 0.013 | 0.478 ± 0.013 | 0.512 ± 0.027 | 0.494 ± 0.049 | 0.512 ± 0.027 | 0.494 ± 0.049 | 0.512 ± 0.027 |
| | AUC | 0.500 ± 0.000 | 0.500 ± 0.000 | 0.492 ± 0.026 | 0.474 ± 0.053 | 0.492 ± 0.027 | 0.474 ± 0.053 | 0.492 ± 0.027 |
| 12 months | Accuracy | — | — | — | 0.806 ± 0.059 | 0.857 ± 0.031 | 0.807 ± 0.058 | **0.869 ± 0.048** |
| | AUC | — | — | — | 0.883 ± 0.056 | 0.912 ± 0.044 | 0.877 ± 0.054 | **0.917 ± 0.045** |
| 30 months | Accuracy | — | — | — | 0.784 ± 0.062 | 0.811 ± 0.055 | 0.788 ± 0.063 | **0.878 ± 0.035** |
| | AUC | — | — | — | 0.842 ± 0.061 | 0.871 ± 0.062 | 0.870 ± 0.046 | **0.933 ± 0.027** |
| 60 months | Accuracy | — | — | — | 0.764 ± 0.056 | 0.759 ± 0.061 | 0.783 ± 0.063 | **0.867 ± 0.030** |
| | AUC | — | — | — | 0.814 ± 0.063 | 0.802 ± 0.051 | 0.874 ± 0.053 | **0.937 ± 0.018** |
| **Permissive profile** |  |  |  |  |  |  |  |  |
| Baseline | Accuracy | 0.430 ± 0.015 | 0.570 ± 0.015 | 0.508 ± 0.049 | 0.524 ± 0.070 | 0.539 ± 0.027 | 0.524 ± 0.070 | 0.511 ± 0.043 |
| | AUC | 0.500 ± 0.000 | 0.500 ± 0.000 | 0.518 ± 0.041 | 0.476 ± 0.057 | 0.543 ± 0.045 | 0.476 ± 0.057 | 0.532 ± 0.065 |
| 12 months | Accuracy | — | — | — | 0.795 ± 0.069 | **0.814 ± 0.049** | 0.788 ± 0.032 | 0.811 ± 0.033 |
| | AUC | — | — | — | 0.874 ± 0.039 | **0.883 ± 0.036** | 0.866 ± 0.035 | 0.875 ± 0.042 |
| 30 months | Accuracy | — | — | — | 0.784 ± 0.040 | 0.785 ± 0.064 | 0.796 ± 0.048 | **0.841 ± 0.040** |
| | AUC | — | — | — | 0.823 ± 0.055 | 0.835 ± 0.070 | 0.856 ± 0.054 | **0.900 ± 0.040** |
| 60 months | Accuracy | — | — | — | 0.722 ± 0.055 | 0.721 ± 0.080 | 0.807 ± 0.050 | **0.837 ± 0.052** |
| | AUC | — | — | — | 0.738 ± 0.057 | 0.755 ± 0.041 | 0.865 ± 0.043 | **0.910 ± 0.037** |

At baseline, the available measurements provided little discrimination between the preferred-treatment classes. Among the learned models, AUC values ranged from 0.474 to 0.492 under the conservative profile and from 0.476 to 0.543 under the permissive profile. The constant strategies yielded AUC values of 0.5 by construction, and the baseline MLP provided similar prediction performance. Thus, although baseline measurements were moderately predictive of the final OFC outcome, they were insufficient to reliably identify the retrospectively preferred treatment.

Preferred-treatment prediction improved markedly when the recommendation was updated using measurements obtained at 12 months. Under the conservative profile, the History-aware MLP achieved the highest mean performance, with an accuracy of 0.869 and an AUC of 0.917. Under the permissive profile, the Visit-wise MLP achieved the highest corresponding values, with an accuracy of 0.814 and an AUC of 0.883. These results show that the first follow-up assessment provided substantial treatment-relevant information that was not available at baseline.

High discrimination was maintained at the subsequent visits, with the greatest later-visit performance generally obtained when the accumulated clinical history was included. Under the conservative profile, the History-aware MLP achieved AUC values of 0.933 and 0.937 at 30 and 60 months, respectively. Under the permissive profile, it achieved AUC values of 0.900 and 0.910 at the same visits. Its corresponding accuracies were 0.878 and 0.867 under the conservative profile and 0.841 and 0.837 under the permissive profile. The benefit of history was therefore most apparent at the later visits, when a longer sequence of clinical and immunological measurements was available.

The tree-based models also benefited from visit-wise reassessment, demonstrating that the information acquired during follow-up could be used by both structured decision-tree models and multilayer perceptrons. However, the History-aware MLP consistently outperformed the History-aware DT at all follow-up visits under both preferred-treatment profiles. This pattern is consistent with the MLP capturing multivariable relationships across the accumulated clinical history that were not represented as effectively by the corresponding tree-based models.

Together, these findings establish that preferred-treatment selection benefits from being formulated as an evolving, visit-wise adaptive problem rather than as a single baseline decision. When baseline information was limited, the recommendation remained uncertain; after follow-up measurements became available, preferred treatment could be predicted with substantially greater accuracy and discrimination, with additional gains at later visits from incorporating the accumulated clinical history. However, these adaptive models optimized each visit independently and could therefore generate stop, restart sequences that are incompatible with irreversible treatment discontinuation. We next examine how imposing this clinical constraint affects the apparent benefit of visit-wise adaptation.

### Irreversible discontinuation erodes the benefit of visit-wise adaptation

The preceding analysis evaluated visit-wise and history-aware models under a fully flexible policy, in which the recommendation at each visit was scored independently of earlier decisions. We next evaluated the same predictions under the clinically relevant constraint that treatment could not be resumed after discontinuation. Once avoidance was predicted, all subsequent recommendations were therefore set to avoidance. Denoting consumption by 1 and avoidance by 0, a predicted trajectory such as (1, 0, 1, 0) was transformed into (1, 0, 0, 0), whereas the admissible trajectory (1, 1, 1, 0) remained unchanged and (0, 1, 1, 0) became (0, 0, 0, 0). The constrained analysis used the same trained models, cross-validation splits, decision thresholds, and out-of-fold prediction scores as the unconstrained analysis. The only difference was the post hoc application of the irreversible-discontinuation rule to the predicted trajectories.

Imposing irreversible discontinuation substantially reduced the performance of the independently optimized adaptive models under both preferred-treatment profiles (Table 3, Fig. 6). Across models and follow-up visits, accuracy typically decreased by approximately 15–25 percentage points, while AUC decreased by approximately 20–30 percentage points. Comparable losses were observed under the conservative and permissive profiles and affected both visit-wise and history-aware models. Thus, access to accumulated clinical measurements improved local prediction but was insufficient to preserve adaptive performance once early discontinuation restricted all subsequent treatment decisions.

**Table 3.** Performance under irreversible-discontinuation constraint. Values show mean accuracy and AUC ± standard deviation across 10 patient-level cross-validation folds. For every model, the first predicted avoidance decision was propagated through all subsequent visits. Visit-wise models use measurements from the corresponding visit, whereas history-aware variants use all measurements accumulated through that visit. The highest mean performance among the models at each follow-up visit is shown in bold; bolding does not indicate statistical significance.

| Visit | Metric | Visit-wise<br>DT | Visit-wise<br>MLP | TreatNet | History-aware<br>DT | History-aware<br>MLP | TreatNet<br>(Hist.) |
| --- | --- | --- | --- | --- | --- | --- | --- |
| **Conservative profile** |  |  |  |  |  |  |  |
| Baseline | Accuracy | 0.494 ± 0.049 | 0.531 ± 0.050 | 0.501 ± 0.031 | 0.511 ± 0.031 | 0.519 ± 0.044 | 0.494 ± 0.031 |
| | AUC | 0.474 ± 0.053 | 0.519 ± 0.052 | 0.529 ± 0.063 | 0.495 ± 0.031 | 0.528 ± 0.070 | 0.507 ± 0.080 |
| 12 months | Accuracy | 0.629 ± 0.112 | 0.607 ± 0.068 | 0.822 ± 0.048 | 0.588 ± 0.064 | 0.663 ± 0.077 | **0.839 ± 0.052** |
| | AUC | 0.618 ± 0.138 | 0.591 ± 0.078 | 0.859 ± 0.051 | 0.570 ± 0.069 | 0.660 ± 0.096 | **0.868 ± 0.063** |
| 30 months | Accuracy | 0.615 ± 0.111 | 0.600 ± 0.074 | 0.828 ± 0.063 | 0.579 ± 0.054 | 0.654 ± 0.073 | **0.834 ± 0.051** |
| | AUC | 0.607 ± 0.121 | 0.589 ± 0.073 | 0.837 ± 0.069 | 0.566 ± 0.057 | 0.651 ± 0.086 | **0.850 ± 0.061** |
| 60 months | Accuracy | 0.618 ± 0.108 | 0.604 ± 0.062 | 0.820 ± 0.056 | 0.585 ± 0.049 | 0.663 ± 0.079 | **0.834 ± 0.055** |
| | AUC | 0.596 ± 0.111 | 0.581 ± 0.064 | 0.824 ± 0.062 | 0.561 ± 0.054 | 0.650 ± 0.085 | **0.841 ± 0.064** |
| **Permissive profile** |  |  |  |  |  |  |  |
| Baseline | Accuracy | 0.471 ± 0.050 | 0.514 ± 0.048 | 0.565 ± 0.025 | 0.467 ± 0.062 | 0.512 ± 0.034 | 0.568 ± 0.016 |
| | AUC | 0.481 ± 0.051 | 0.550 ± 0.041 | 0.433 ± 0.071 | 0.489 ± 0.035 | 0.541 ± 0.034 | 0.441 ± 0.058 |
| 12 months | Accuracy | 0.564 ± 0.117 | 0.589 ± 0.110 | 0.811 ± 0.045 | 0.561 ± 0.149 | 0.610 ± 0.079 | **0.819 ± 0.053** |
| | AUC | 0.608 ± 0.097 | 0.642 ± 0.110 | 0.865 ± 0.043 | 0.614 ± 0.135 | 0.658 ± 0.071 | **0.869 ± 0.047** |
| 30 months | Accuracy | 0.552 ± 0.124 | 0.591 ± 0.128 | **0.819 ± 0.040** | 0.559 ± 0.147 | 0.605 ± 0.074 | 0.812 ± 0.034 |
| | AUC | 0.607 ± 0.107 | 0.638 ± 0.107 | 0.853 ± 0.038 | 0.619 ± 0.130 | 0.657 ± 0.068 | **0.858 ± 0.039** |
| 60 months | Accuracy | 0.529 ± 0.103 | 0.575 ± 0.120 | **0.810 ± 0.046** | 0.560 ± 0.137 | 0.603 ± 0.077 | 0.805 ± 0.034 |
| | AUC | 0.577 ± 0.080 | 0.617 ± 0.099 | 0.819 ± 0.045 | 0.617 ± 0.125 | 0.650 ± 0.065 | **0.843 ± 0.046** |

**Fig 6.**
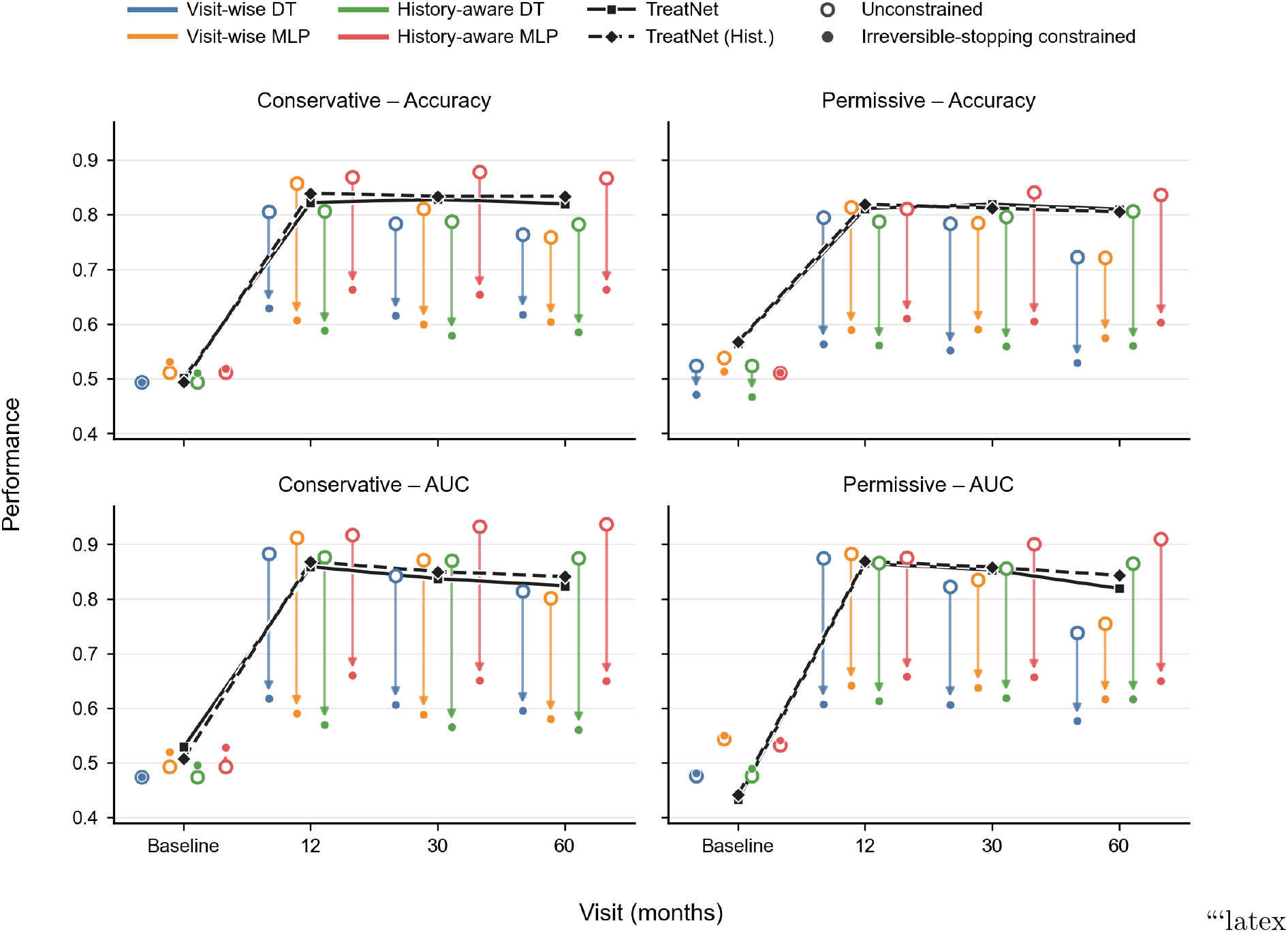
Performance erosion under irreversible discontinuation and comparison with TreatNet. Accuracy and AUC are shown as functions of visit under the conservative and permissive preferred-treatment profiles. For each independently optimized visit-wise or history-aware model, connected points show performance before and after imposing irreversible discontinuation. Under the unconstrained policy, each visit-specific recommendation was evaluated independently; under the constrained policy, the first predicted avoidance decision was propagated through all subsequent visits. The connecting arrows therefore show the performance erosion caused by converting independent predictions into clinically admissible treatment trajectories. Black lines show the performance of TreatNet under the same irreversible-discontinuation constraint. Whereas the independently optimized models lost much of the benefit of longitudinal reassessment after the constraint was imposed, TreatNet preserved higher and comparatively stable performance across follow-up visits.

The unconstrained scores accurately describe the models’ ability to predict the preferred treatment at individual visits, but they do not capture the consequences of converting these predictions into a realizable treatment trajectory. Under independent evaluation, a correct recommendation at a later visit can contribute to performance even when treatment was already discontinued at an earlier visit. Under the irreversible policy, that later recommendation is no longer actionable: an erroneous early discontinuation necessarily propagates through the remainder of the trajectory.

Providing the models with the accumulated patient history did not prevent this degradation. Historical information improved local prediction, particularly at the later visits, but the independently optimized objectives did not account for how an early decision restricts all subsequent treatment options. The limitation therefore lies not only in the information provided to the model, but also in the mismatch between visit-level optimization and the path-dependent structure of the clinical decision.

These results distinguish predictive adaptivity from clinically coherent adaptivity. Reassessing treatment over time provides a substantial advantage over fixing the decision at baseline, but this advantage is not preserved when an irreversible constraint is imposed only after training. Preserving the value of longitudinal reassessment requires an objective that accounts for the downstream consequences of each discontinuation decision while the model is being optimized. This provides the motivation for the trajectory-aware TreatNet formulation.

### TreatNet preserves adaptive performance under irreversible discontinuation

Since the constraint of irreversible discontinuation erodes the benefit of independently optimized adaptive models, we next examined whether incorporating this path dependence during training could preserve the value of longitudinal reassessment. TreatNet provides a controlled comparison with the Visit-wise MLP: both use the same visit-specific architecture and receive the same clinical information, but TreatNet couples the visit-specific outputs through a trajectory-level objective that accounts for the downstream consequences of early discontinuation. The same irreversible stopping rule was applied to TreatNet and all reference models during evaluation.

Across all follow-up visits and both preferred-treatment profiles, TreatNet achieved substantially higher mean performance than the independently optimized adaptive models under the same irreversible-discontinuation constraint (Table 3, Fig. 6). The TreatNet variants attained accuracies of 0.805–0.839 and AUC values of 0.819–0.869, whereas the reference models did not exceed an accuracy of 0.663 or an AUC of 0.660. The advantage was already evident at 12 months: under the conservative profile, History-aware TreatNet achieved an accuracy of 0.839 and an AUC of 0.868, compared with 0.663 and 0.660, respectively, for the strongest non-TreatNet comparators. Under the permissive profile, the corresponding TreatNet values were 0.819 and 0.869, whereas the strongest reference models reached an accuracy of 0.610 and an AUC of 0.658. Comparable performance was maintained at 30 and 60 months under both profiles, indicating that trajectory-aware training preserved the benefit acquired at the first follow-up assessment throughout the remaining treatment pathway.

Visit-wise reassessment revealed preferred-treatment information that was largely unavailable at baseline, but independently optimized models could not retain this advantage after their predictions were converted into irreversible treatment trajectories. TreatNet preserved the benefit because the consequences of early discontinuation were represented during optimization rather than imposed only after prediction.

It is important to note that the improvement did not result from access to additional clinical information or future measurements. At inference, each TreatNet module used only the information available at its corresponding visit, exactly as in the matched MLP reference model. The difference lay in the training objective: coupling the visit-specific outputs allowed prediction errors to be weighted according to their consequences for the remaining treatment pathway. Early discontinuation was therefore optimized as a trajectory-level decision rather than as an isolated visit-level classification.

Together, these results define a progression from static treatment selection to visit-wise adaptation and, finally, to clinically coherent adaptive decision-making. Reassessment over time first exposed treatment-relevant information that was unavailable at baseline. Irreversible discontinuation then showed that this opportunity could not be realized reliably through independent visit-level optimization. By incorporating path dependence during training, TreatNet retained high predictive performance while producing recommendations compatible with the admissible treatment pathway. Thus, when treatment discontinuation is irreversible, the value of longitudinal information depends not only on when predictions are updated, but also on whether the learning objective reflects the downstream consequences of acting on them.

### TreatNet defers and selectively applies treatment discontinuation

To characterize the strategy underlying TreatNet’s performance, we examined how History-aware TreatNet reclassified patients between baseline and 12 months (Figure 7). Out-of-fold predictions from the 10 outer cross-validation folds were pooled from the LEAP data.

**Fig 7.**
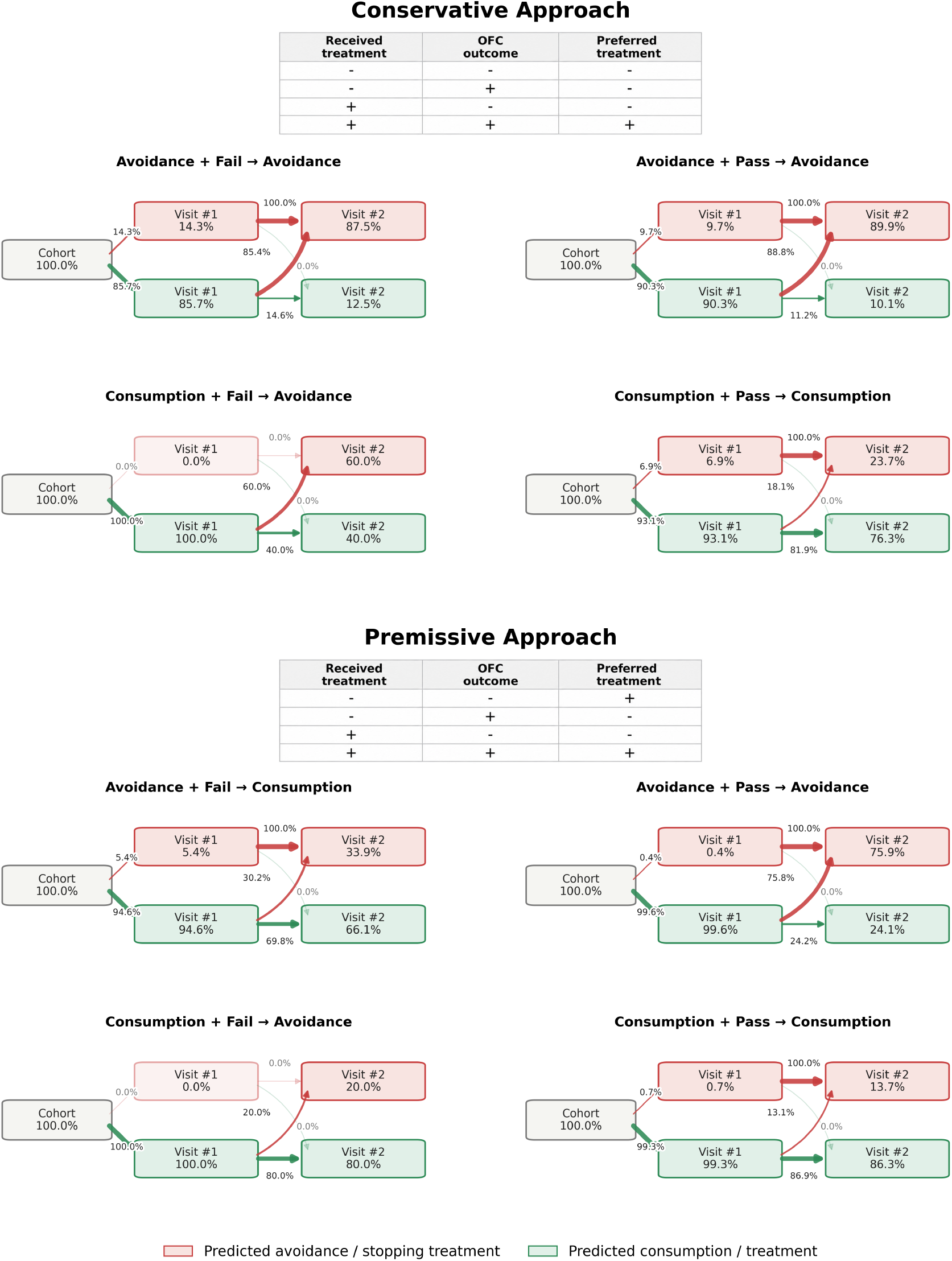
TreatNet defers and selectively applies irreversible discontinuation as longitudinal information becomes available. Flow diagrams show predictions from History-aware TreatNet at baseline and 12 months under the conservative and permissive preferred-treatment profiles. Percentages within each visit block show the distribution of predicted consumption and avoidance for that subgroup, whereas percentages on the connecting arrows show the conditional transitions between baseline and 12 months. Because discontinuation is irreversible, patients predicted to avoid at baseline cannot subsequently return to consumption. Across profiles, TreatNet generally preserves consumption while baseline information is limited and applies selective, profile-dependent discontinuation after the more informative 12-month measurements become available. The profiles differ for patients assigned to avoidance who failed the final OFC: the conservative profile defines avoidance as their preferred treatment, whereas the permissive profile defines consumption as preferred. Accordingly, TreatNet adopts a more avoidance-oriented strategy under the conservative profile and a more consumption-oriented strategy under the permissive profile.

Across both preferred-treatment profiles, TreatNet predominantly recommended consumption at baseline, including for many patients whose retrospective label was avoidance. This behavior reflects the asymmetry of irreversible discontinuation: continuing consumption preserves the opportunity for later reassessment, whereas stopping treatment removes that option. After the more informative 12-month measurements became available, recommendations shifted toward avoidance primarily in subgroups for which avoidance was the retrospective preferred-treatment label.

Under the conservative profile, the proportion predicted to avoid increased from 14.3% to 87.5% among avoidance/OFC-fail patients and from 9.7% to 89.9% among avoidance/OFC-pass patients. Among consumption/OFC-fail patients, all five were initially predicted to consume, and 60.0% were redirected to avoidance at 12 months. By contrast, 93.1% of consumption/OFC-pass patients were initially predicted to consume, and 76.3% retained that recommendation at follow-up. Discontinuation was therefore concentrated in subgroups labeled as preferring avoidance rather than applied uniformly across the cohort.

The transition pattern changed under the permissive profile, in which avoidance/OFC-fail patients were labeled as preferring consumption. TreatNet recommended consumption to 94.6% of this subgroup at baseline and retained consumption for 66.1% at 12 months, compared with only 12.5% under the conservative profile. For avoidance/OFC-pass patients, whose preferred-treatment label remained avoidance, the proportion predicted to avoid increased from 0.4% to 75.9%. Consumption was retained at 12 months for 86.3% of consumption/OFC-pass patients, whereas only one of the five consumption/OFC-fail patients was redirected to avoidance.

The largest clinically relevant transition occurred among the 257 avoidance/OFC-pass patients, who were labeled as preferring avoidance under both profiles. TreatNet initially preserved consumption for most of these patients and discontinued it only after the 12-month assessment, identifying the principal opportunity for reducing potentially unnecessary treatment exposure. Reclassification of the five consumption/OFC-fail patients may represent a second clinically relevant pattern, but the small subgroup size makes the corresponding percentages descriptive rather than conclusive.

Some consumption/OFC-pass patients also transitioned to avoidance, despite consumption being their retrospective preferred-treatment label. These transitions are errors relative to the constructed label, but their clinical meaning remains uncertain because outcomes under earlier discontinuation were not observed. The analysis therefore characterizes the trade-off learned by TreatNet under the retrospective labels rather than establishing the safety or optimality of discontinuation for individual patients.

Overall, TreatNet learned a defer-then-discontinue strategy: it generally preserved consumption when baseline information was limited and applied irreversible discontinuation selectively after the 12-month assessment. The marked difference between the conservative and permissive profiles shows that this behavior was not a fixed tendency to stop treatment. Instead, the continuation threshold adapted to the preferred-treatment definition, demonstrating how trajectory-aware optimization integrates uncertainty, accumulating evidence, and the consequences of irreversible action into a coherent longitudinal decision policy.

## Discussion

Training a decision-support system for longitudinal treatment selection is challenging because the model must learn when the available evidence is sufficient to support a treatment change. This problem becomes particularly difficult when treatment discontinuation is irreversible, because an early decision to stop treatment constrains all subsequent decisions and prevents later information from being acted upon. Such a constraint is relevant to peanut-allergy prevention, where resuming peanut consumption after discontinuation may require renewed clinical assessment. In this work, we introduce a longitudinal framework for preferred-treatment prediction under irreversible discontinuation and evaluate it using data from children assigned to early peanut consumption or avoidance in the LEAP trial. We first introduce visit-wise adaptive and history-aware models and show that preferred treatment, although difficult to determine at baseline, becomes substantially more predictable as follow-up information accumulates. We then show that much of this advantage is lost when irreversible discontinuation is imposed on models optimized independently at each visit. To address this limitation, we introduce TreatNet, a trajectory-aware model that incorporates the downstream consequences of discontinuation into training, thereby preserving the value of longitudinal reassessment while producing treatment trajectories that satisfy the irreversible-discontinuation constraint.

The longitudinal analyses showed corresponding changes in OFC predictive performance and clinical feature importance (Fig. 5). Baseline measurements provided only moderate discrimination of the final OFC outcome and limited discrimination of the retrospectively preferred treatment. Predictive performance increased as follow-up information became available, while the features contributing most strongly to OFC prediction also changed. Treatment assignment was particularly informative at baseline, whereas physiological and immunological measurements, including skin-prick-test wheal size and the IgG4/IgE ratio, became more prominent at later visits. Thus, the information available for outcome prediction was not static: the relative importance of treatment exposure, clinical response, and immunological state changed over time. A recommendation made only at baseline cannot capture this evolving clinical signal or use information subsequently acquired through longitudinal monitoring.

Consistent with this evolution, models that reassessed preferred treatment during follow-up substantially outperformed strategies that retained a single baseline recommendation (Table 2). The largest improvement occurred after the first follow-up assessment, when the available measurements better distinguished treatment preferences that were poorly resolved at baseline. History-aware models, which incorporated all measurements accumulated through each visit, generally provided an additional benefit at later visits. This improvement did not depend on observing the final outcome or accessing future measurements: every prediction used only the information available at the corresponding clinical time point. These results show that longitudinal reassessment provides an opportunity to revise an initially uncertain recommendation as the child’s clinical and immunological trajectory becomes clearer. They also distinguish longitudinal treatment selection from baseline prediction, in which a single recommendation remains fixed throughout follow-up.

Accurate prediction at individual visits, however, does not necessarily produce a coherent treatment trajectory. This distinction becomes critical when discontinuation changes which actions remain available. Under an irreversible policy, an erroneous early decision to discontinue cannot be corrected later, even if subsequent measurements favor continued treatment. Errors are therefore temporally asymmetric: a late error affects a limited portion of the trajectory, whereas an early discontinuation propagates through every remaining visit. Discontinuation consequently cannot be represented adequately as a post-processing rule applied after independent predictions have been generated. We demonstrate this mismatch through the substantial erosion in accuracy and discrimination observed when the irreversible constraint was imposed on the visit-wise adaptive models (Table 3, Fig. 6). History-aware models showed similar degradation, indicating that access to accumulated information cannot compensate for a training objective that disregards the consequences of acting on an early prediction.

TreatNet uses visit-specific predictors and only the clinical information available at each visit. Rather than optimizing each predictor independently, however, TreatNet couples their outputs through a trajectory-level objective. An early discontinuation is therefore evaluated according to its effect on the remaining treatment pathway rather than as an isolated classification at a single visit. This formulation allows the model to learn both which treatment is preferred at a given assessment and how the timing of an irreversible decision affects the complete trajectory (Table 3, Fig. 6). TreatNet does not derive its advantage from additional variables, future information, or a more expressive visit-level architecture. Its technical novelty lies in aligning model optimization with the path-dependent structure of the decision problem. Comparison with the independently trained MLP models therefore isolates the value of trajectory-aware optimization from that of the underlying neural predictors.

When evaluated under irreversible discontinuation, TreatNet generally achieved stronger discrimination than the independently optimized adaptive models while maintaining competitive accuracy across follow-up visits and both preferred-treatment profiles. The transition analysis provides an interpretable account of this behavior. When baseline evidence was limited, TreatNet generally preserved peanut consumption, thereby retaining the opportunity to reassess treatment later (Figure 7). After the more informative 12-month assessment, it selectively shifted patients toward avoidance, particularly in subgroups for which avoidance was retrospectively preferred. This defer-then-discontinue pattern reflects the asymmetry of the decision: continuation preserves future options, whereas premature discontinuation removes them. TreatNet therefore used information gained during follow-up not simply to update predictions, but also to learn the timing of irreversible discontinuation.

The transition patterns also differed between the conservative and permissive preferred-treatment profiles (Figure 7). Under the conservative profile, TreatNet favored discontinuation more strongly for patients assigned to avoidance who failed the final OFC. Under the permissive profile, in which continued consumption was preferred for this subgroup, the model retained consumption more frequently. TreatNet therefore did not learn a fixed tendency to continue or discontinue treatment; its stopping behavior adapted to the preferred-treatment definition used during training. More broadly, this comparison illustrates why predicting a clinical outcome is not equivalent to selecting a treatment. Treatment selection additionally requires assumptions about the relative consequences of treatment failure, unnecessary exposure, premature discontinuation, and continued treatment burden. The two profiles were used to examine the effect of these assumptions rather than to assert a single definitive clinical preference.

Prospective evaluation in independent and more diverse cohorts will be needed to establish the clinical utility, safety, and transportability of TreatNet. Beyond this necessary validation, the framework addresses a general class of longitudinal decision problems in which an action changes the options available later. Although evaluated here for early peanut consumption, TreatNet is not specific to peanut-allergy prevention or binary treatment discontinuation. Its trajectory-level formulation can represent irreversible or difficult-to-reverse decisions, multiple treatment states, asymmetric transition costs, minimum treatment durations, treatment switching, and escalation or de-escalation pathways. The same principle is relevant to medication tapering, preventive therapy, rehabilitation, and chronic-disease management, where optimizing each assessment independently may produce a sequence of predictions that is incompatible with the actual treatment pathway. TreatNet provides a general approach for aligning longitudinal prediction with these pathway constraints, allowing decision-support models to optimize clinically realizable trajectories rather than isolated recommendations.

## Acknowledgment

This work was supported by the Food Allergy Fund (FAF), the Israeli Ministry of Science and Technology (MOST) grant #7195, and the Zimin Institute for AI in Healthcare.

